# Two-photon microscopy at >500 volumes/second

**DOI:** 10.1101/2020.10.21.349712

**Authors:** Yu-Hsuan Tsai, Chih-Wei Liu, Wei-Kuan Lin, Chien-Sheng Wang, Chi-Huan Chiang, Vijay Raj Singh, Peter T. C. So, Chia-Fu Chou, Shi-Wei Chu

## Abstract

We demonstrate a multi-focal multi-photon volumetric microscopy via combination of 32-beam parallel lateral-scanning, a 70-kHz axial-scanning acoustic lens, and a 32-channel photodetector, enabling unprecedented data rate (2-10 GHz) and >500-volumes/second imaging speed over ~200×200×200-μm^3^.

---

Recently, one major trend of optical microscopy is to improve volumetric imaging speed toward millisecond-scale, which is highly desirable for rapid transient events, such as neural activity and flow cytometry. Light sheet technique allows hundreds of volumes per second [1], but its wide-field detection degrade quickly with scattering in deep-tissue imaging. To enhance imaging depth, two-photon volumetric microscopy combining various fast axial imaging schemes have been demonstrated [2–7]. The first three exhibit imaging speeds at only ~10 volumes per second because their methods require sequentially raster scanning frame by frame [2–4]. Bessel-beam-based technique [5] reaches 50 Hz volumetric imaging rate, but limited to sparse samples. The last two methods achieve millisecond-scale sampling of multiple points in 3D, but is susceptible to motion artifact [6,7]. The state-of-the-art multi-photon microscopy can reach thousands of frames per second [8], but limited to 2D observation. Note that the abovementioned two-photon techniques use a single-channel detector, so due to limitation of fluorescence lifetime (2-3 ns), the data acquisition rate is restricted below 500 MHz. To further enhance imaging speed, one strategy is to introduce parallel scanning [9] or wide-field excitation [10]. However, those schemes are mostly based on wide-field detection, which again faces the challenge of limited penetration depth. The design may be further improved via adding multiple point detectors [11], which offer much better penetration depth than wide-field camera detectors.

In this study, we developed a MUlti-focal MUlti-photon Volumetric Imaging Microscopy (MUVIM), based on combination of 32-beam parallel scanning by a diffractive optical element (DOE), ~70 kHz ultrafast axial scanning by a tunable acoustic gradient-index (TAG) lens [12], and a 32-channel photomultiplier tube (PMT) that boosts up data acquisition rate to more than 10 GHz (2.56 GHz is used in this work), thus reaching unprecedented volumetric imaging rate above 500 volumes per second, in a cubic volume of ~200 μm on each side. Our record high-speed multi-photon volumetric imaging paves the way toward applications of high-throughput flow cytometry and functional neural network analysis, for examples.

The overall setup is shown in Fig. 1(a). An 80-MHz Ti:Sapphire laser was sent through a DOE to form 32 collimated beams (Fig. 1(b)), which were then relayed onto a galvanometric mirror for lateral scanning (Fig. 1(c)). These scanning beams were projected onto the back aperture of a TAG lens (TAG Lens 2.5*β*, TAG Optics Inc.), to scan the 32 foci simultaneously up and down at 70 kHz, forming multiple ribbon scan (Fig. 1(d) [12]. After an objective lens (HC PL IRAPO 20x, NA=0.75, Leica), a volume of ~200 × 220 × 170 μm^3^ was covered within 1.84 millisecond(maximal volume size 200(x) × 800(y) × 200(z) μm^3^). Fig. 1(e) and 1(f) show the lateral and axial resolution as 690 nm and 5.5 μm, respectively, both close to theoretical estimation (630 nm and 4.6 μm). The slight deviation may be due to intrinsic system aberration. The distance among each focus after objective was set at 6.7 μm, which could be tuned by different lens combination. Maximum laser power at focal plane was typically 120 mW. The emitted fluorescent signal (80 MHz) was epi-detected by a 32-channel linear PMT (H12175-200, Hamamatsu), resulting in signal bandwidth at 2.56 GHz.

**Fig. 1.**
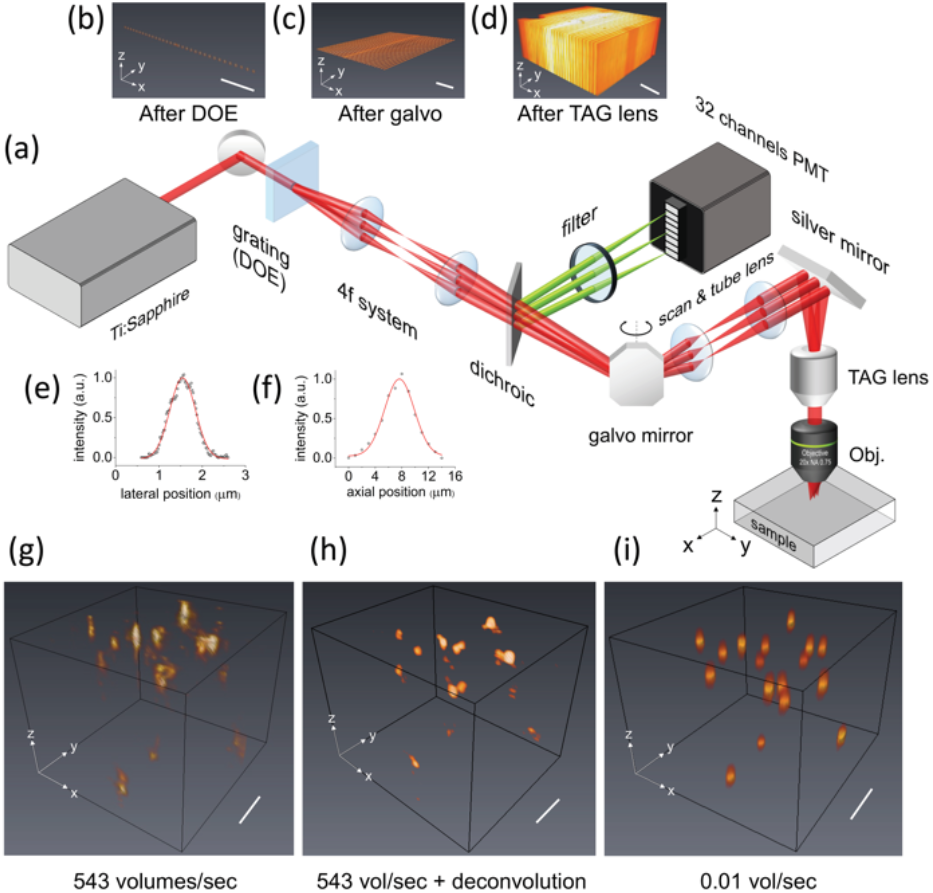
(a) Schematic of MUVIM. DOE, diffractive optical element; TAG, tunable acoustic gradient-index; Obj., objective; PMT, photomultiplier tube; galvo, galvanometric. (b) Image of 32 foci after DOE; (c) Image of 32 scanning lines after galvo; (d) Volumetric image after the TAG lens, acquired with a thin green-fluorescent film and a camera. (e)(f) Lateral and axial resolution respectively. (g) 3D Observation of 10-μm fluorescent microspheres (1.84 millisecond) by MUVIM, (h) by adding deconvolution, and (i) by slow single beam scanning. All scale bars are 50 μm.

Fig. 1(g) shows one representative result of ultrahigh speed volumetric imaging of 10 μm fluorescent microspheres (see Visualization 1 for dynamics with more than 500 volumes per second). To our knowledge, this is the highest two-photon volumetric imaging rate to date. The location of each microsphere was verified througha single beam raster scanning in Fig. 1(i), which took a few minutes to obtain one volume. The high-speed imaging is based on trade-off of lower lateral sampling density (32 × 128 in Fig. 1(g) versus 512 × 512 in Fig. 1(h)) and reduced signal-to-noise ratio. Nevertheless, the location of each microsphere was precisely mapped, manifesting the potential toward studying dynamics in neuronal system.

For high-speed imaging, data acquisition rate is one key factor since it determines the upper limit of imaging speed. In our system, the maximal data throughput of each PMT channel reaches more than 300 MHz, so the 32-channel PMT leads to more than 10 GHz (now limited to 2.56 GHz by the laser pulse repetition rate), similar to the state-of-the-art parallel camera-based methods [13]. Nevertheless, theframerateofcameracannotcatchup with the TAG lens, highlighting the necessity of multi-channel PMT.

Another major advantage of multi-channel PMT in a multifocal multiphoton system over camera detection is to eliminate crosstalk in scattering tissues, through photon reassignment and deconvolution, thus enhancing imaging depth [11][14]. Fig. 1(h) presents 3D deconvolved image, as a proof of concept that our system is ready for deep-tissue volumetric high-speed imaging.

The high-speed volumetric imaging opens the possibility of not only 3D complex cellular structure monitoring, such as an organoid [15], in flow cytometry, but also dynamic observation in brain neuroscience [16]. Our two-photon volumetric imaging system may allow high-speed organoid screening with sub-cellular precision. A quick estimation shows that it is possible to count nearly 4 million cells per second in our system (assuming cells in organoids are 10-μm tightly packed spheres, i.e. ~ 20^3^ cells in 200 × 200 × 200 μm^3^ volume, which is imaged within 2 ms), suggesting the potential of “flow organoidmetry”. Regarding neuroscience, one important model animal is *Drosophila*, whose brain size is on the order of 200 × 400 × 200 μm^3^. Our system is ready toward *in vivo* whole-brain functional mapping, i.e. capturing millisecond dynamics of action potential communications among neurons in a living brain. The main challenge is photon yield under high speed sampling, in particular for fluorescent proteins. For example, under 1 μM concentration of eGFP, the estimated fluorescent photon yield is ~ 100 photons per laser pulse. The strategies to improve the photon yield include increasing labeling concentration, laser power, repetition rate, reducing pulse width, etc. These will be our future research direction.

In summary, our MUVIM system achieves record-high 500 volumes per second with sub-micrometer focal spot size in a ~200 μm cubic volume, based on the combination of not only the high-speed sampling of multi-focus laser scanning and the TAG lens, but also the deep imaging capability of two-photon excitation plus multiple point detectors, which enables >10 GHz data throughput. This innovative high-speed volumetric imaging system may open the avenue for high-throughput cytometry and millisecond-scale neural dynamics study in living brains.

